# Droplet-array culture enables single-cell isolation of bacteria directly from blood

**DOI:** 10.64898/2026.10.15.759347

**Authors:** Alexander Molenda, Marcos Lares, Nathan Hendrikse, Brittney Hynes, Sue McCrone, Warren E. Rose, David J. Beebe

**Affiliations:** Department of Biomedical Engineering, University of Wisconsin-Madison, Madison, WI, USA; School of Medicine and Public Health, University of Wisconsin-Madison, Madison, WI, USA; Carbone Cancer Center, University of Wisconsin-Madison, Madison, WI, USA; University of Wisconsin Hospitals and Clinics, Clinical Laboratories, Madison, WI, USA; School of Pharmacy, University of Wisconsin-Madison, Madison, WI, USA; Department of Pathology and Laboratory Medicine

## Abstract

Bacteremia and sepsis require rapid diagnosis and treatment, but these are often hampered by slow or inconclusive blood culture results. Here we describe an alternative to traditional blood culture, leveraging common laboratory equipment and a droplet array to isolate and expand bacteria in a single step. The method neutralizes inhibitory whole-blood components through lysis, concentrates sample bacteria, and uses a droplet array to enable single-bacteria isolation and subsequent in-droplet expansion. Seven species of bacteria (four gram negative, three gram positive) were spiked into 1 mL of whole blood at a concentration of approximately 100 colony-forming units (CFU) and subsequently processed. The mean time to colorimetric detection via WST-8 dye was 11.5 h (range 9-13 h), compared with ≥ 28 h typically required for conventional blood culture and subculture. Colony counts per droplet followed a Poisson distribution for both gram-negative (*λ* = 0.288) and gram-positive (*λ* = 0.320) organisms, and means of 86.2% and 88.4% of occupied droplets, respectively, contained a single CFU. This method eliminates the need for additional subculture after positive blood culture. By partitioning the sample, the workflow limits competitive exclusion and preserves the composition of polymicrobial samples, while yielding viable colonies substantially faster than conventional methods. Blood processing at low CFU counts (3-5 CFU/mL) with both *S. aureus* and *E. coli* consistently recovered viable bacteria (1-3 CFU/mL).

**Importance:** Bloodstream infections require prompt, appropriate antibiotic therapy, but current laboratory workflows responsible for identifying pathogens can take 24 hours at minimum. Blood cultures require bacteria to grow to detectable levels in the presence of immune cells and other inhibitory blood components, followed by an additional overnight subculture to obtain the individual colonies required for identification and subsequent susceptibility testing. We describe a method that shortens the culture process by removing inhibitory blood components, concentrating any bacteria present, and rapidly distributing them across an array of droplets where growth and isolation occur concurrently. Viable colonies for downstream testing were recovered in approximately 11 hours using standard laboratory equipment and inexpensive reagents. This method of partitioning bacteria before expansion may also preserve minority populations that may be excluded or outcompeted in a single bulk culture.

## Introduction

Sepsis, a dysregulated host immune response to microbial infection, can result in major organ failure and death. Sepsis constitutes a significant global health burden, representing 20% of all global deaths. Up to 40% of sepsis cases occurred in children under the age of 5, with areas of low socioeconomic status suffering higher mortality due to limited healthcare resources.^1^ Time is one of the most critical components in the treatment of bacteremia and sepsis, with a risk-adjusted mortality increase of 3-9% for every hour that antibiotic administration is delayed, though this varies with illness severity.^2–5^ While the exact percent increase remains debated, it is understood that prompt treatment helps to improve patient outcomes. However, treatment urgency must be balanced with appropriate antibiotic coverage, as both inadequate and unnecessary antibiotic treatment are associated with higher mortality and contribute to the development of antimicrobial resistance (AMR).^6,7^ Conventional blood culture and susceptibility testing can take days to yield actionable results, delaying the transition from empiric to targeted therapy. During this time, inter and intraspecies competitive exclusion can take place within the blood cultures, causing one or more microbes within a bacterial population to dominate the culture while other strain populations are lost entirely.^8,9^ This bacterial exclusion can confound the true composition of the infection. Diagnostic difficulties or delays force physicians to treat patients empirically with antibiotics, attempting to deliver prompt treatment while hitting a narrow therapeutic target. Any inadequate treatment contributes to the broader burden of AMR. This is especially apparent in cases of heteroresistance, where a majority susceptible population of bacteria contains a small resistant subpopulation that is easily outcompeted due to the fitness cost of resistance genes.^10^ Thus, there is a critical need for a more efficient process for blood culture and bacterial identification that can provide clinicians with faster, more complete information; particularly in cases of polymicrobial infection, AMR, and heteroresistance.

Current gold standard laboratory practices for identifying these pathogens consists of blood culture, followed by agar subculture and identification methods such as matrix-assisted laser desorption/ionization - time of flight (MALDI-TOF), polymerase chain reaction (PCR), 16S rRNA sequencing, or whole genome sequencing (WGS).^11^ Blood culture requires drawing patient blood directly into blood culture bottles and incubating in whole-blood batch culture, typically requiring 12-48 hours for common pathogens, with more fastidious organisms requiring additional time.^12^ During whole blood incubation, bacteria need to overcome the presence of immune cells and other inhibitory components of whole blood while replicating to a detectable density before flagging positive via internal fluorescent CO_2_ sensors. The positive culture must then be subcultured on agar for 4-16 hours to isolate single colonies. Several isolation methods have been proposed to improve the blood culture and bacterial identification process, including filtration, centrifugation, functionalized magnetic beads, and microfluidic chips.^13^ Despite functionally being able to isolate bacteria from whole blood, these techniques all suffer from poor compatibility within clinical microbiology processes, with limitations such as dependence on pre-culture, size-based exclusion, and high technical complexity.

Here we describe a simple workflow capable of isolation and detection of common bacterial pathogens directly from whole blood, using common laboratory infrastructure, inexpensive reagents, and a colorimetric indicator for droplets containing bacteria. The described process produces viable bacteria in 11.5 hours on average after processing (range 9-13 h) by neutralizing whole blood components through sequential ammonium chloride and osmolysis. By preserving and concentrating existing bacteria within the sample, blood culture/subculture steps can be combined using an under-oil droplet array.^14,15^ Compared to the ≥28 hours required for traditional blood culture and agar subculture, this represents a clinically meaningful reduction of 15-19 hours. The lysing of leukocytes and other components of whole blood and early partitioning of the sample helps preserve sample bacterial populations.

In this study we demonstrate that the droplet array can reliably isolate 1 colony-forming unit (CFU) per droplet at clinically relevant bacterial loads (∼100 CFU/mL), producing viable subcultured bacteria in a single culture step.^16,17^ The partitioning method also enables potential identification of heteroresistance and polymicrobial infection, while helping prevent sample-wide contamination.

## Materials and Methods

### Preparation of the Under-oil Droplet Array

A more detailed protocol can be found in previous publications, with the process consisting of ELR surface treatment, surface patterning, and sample seeding.^14^ The glass substrate was prepared with oxygen plasma for 3 min at 100 W (Diener Electronic Femto, Plasma Surface Technology), causing surface activation and allowing for modification with a monolayer of covalently bonded PDMS-silane (1,3-Dichloro-1,1,3,3-tetramethyldisiloxane, TCI, D2334) molecules applied utilizing vapor diffusion. A pattern was then transferred to the prepared glass surface using a PDMS or rubber stamp by an additional oxygen plasma treatment for 1 min at 100 W, creating a hydrophilic pattern on a superhydrophobic surface.

### Bacterial preparation and inoculation

All bacterial strains were sourced from ATCC or as cited (see Supplementary Table 1) and stored at -80 °C in 10% glycerol. Bacteria was plated on Mueller-Hinton agar (Millipore Sigma Mueller Hinton Agar, Catalog No. 70191) and incubated overnight at 37 ºC prior to use. A small portion of bacteria were moved to a glass test tube containing Mueller-Hinton broth (MHB) (BD BBL Dehydrated Culture Media: Mueller Hinton II Broth (Cation-Adjusted), Catalog No. 212322) using a cotton swab, which was then adjusted to McFarland Standard 0.5, or roughly 1.5 × 10^8^ CFU/mL. A serial dilution was performed such that 10 µL was added per milliliter of final solution, producing a final bacterial concentration target of approximately 100 CFU/mL.

### Whole blood processing

*To isolate bacteria, Method 1 (hypotonic osmolysis) was initially used (graphs in fig. 3 and 4, Supplementary Fig. 1). Method 2 replaced hypotonic osmolysis steps with ammonium chloride-based red blood cell lysis to improve visual clarity of positive under-oil droplets using WST-8 for detection*.

Human peripheral blood was drawn into EDTA Vacutainer (BD, GTIN: 00382903666430) by venipuncture from healthy donors with written informed consent under an approved UW Madison Institutional Review Board protocol (2020-1623). Aliquots of blood were added to 1.5mL Eppendorf tubes at 1mL each. Blood was inoculated with bacteria at a desired concentration within 4 hours of blood draw and centrifuged at 20,000 x g for 5 min to separate plasma. Plasma supernatant was aspirated and a washing agent was used to resuspend the remaining pelleted volume of blood. Method 1 utilized 1 mL water (Cytiva HyClone Water, Cell Culture Grade (Endotoxin Free), Catalog no. SH30529). Method 2 used 1 mL 1x red blood cell (RBC) lysis buffer (Miltenyi Biotec Red Blood Cell Lysis Solution (10x), Catalog No. 130-094-183). The sample was vortexed for 5 seconds and incubated at room temperature (RT) for 10 min, followed by centrifugation at 20,000 x g for 2 min. The supernatant was aspirated down to the visible white pellet and an additional 1 mL of washing agent was added to resuspend the pellet containing the bacteria, following the same subsequent vortex, incubation, and centrifugation steps. After centrifugation, the supernatant was removed and water was added up to 1mL as the final wash for both methods. The sample was vortexed for 5 seconds or until the pellet was resuspended, and incubated at RT for 10 min. The sample was centrifuged at 20,000 x g for 2 min, and all but 50 µL of supernatant was removed. WST-8 (20x) (Dojindo, Catalog no. M439-01) was combined with MHB (1:20 dilution) and added to resuspend the concentrated bacterial pellet to a final volume of ∼100 µL. Samples were distributed into an under-oil droplet array.

### Under-oil droplet array seeding and imaging

3 mL of silicone oil (5 cSt) was added to the device, with the concentrated bacterial sample loaded under-oil by standard pipetting or under-oil sweep distribution (fig. 2b,c).^14^ Method 1 primarily utilized sweep distribution. Time lapse imaging was performed on a Nikon Eclipse Ti (10x objective, 10 min interval for 16 h) with an on-stage incubator kept at 37 ºC, 5% CO_2_, and 95% relative humidity (Bold Line, Okolab) (Supplementary Fig. 2). For time lapse imaging experiments, the silicone oil was displaced by Fluorinert FC-40 oil (Sigma-Aldrich Fluorinert FC-40, Catalog no. F9755) and removed to further decrease evaporation.

While optimizing method 2, it was determined that the system could be more readily adopted by unfamiliar users by using standard pipetting and a guide under the device, showing the location of the hydrophilic pattern (fig. 2c). Macroscopic imaging of the color change provided by the WST-8 dye was done over 20 hours within an incubator with a clear window kept at 37 ºC and high relative humidity, Sony α7R III camera with a Sigma 70mm 1:2.8 DG macro lens, and intervalometer (fig. 5a,b, 6a,b).

### Data Analysis and figure plotting

Data analysis and plots were done using Microsoft Excel, GraphPad Prism, and Fiji. Schematics and illustrations were created using BioRender. Code for image segmentation and color pixel thresholding were written in python via Positron with the help of Claude Code by Anthropic.

## Results

### Exclusive liquid repellency (ELR) based underoil droplet array enables rapid isolation and expansion of bacteria

As previously stated, current methodologies for expanding and isolating bacteria from whole blood in a clinical setting rely heavily on blood culture and subsequent agar plate subculture (fig. 1a). Here, we leverage a unique technology called exclusive liquid repellency, or ELR, to combine the steps of blood culture and agar subculture, significantly reducing the time before bacterial identification (fig. 1b). ELR is a repellant treatment that can be applied to various surfaces. When combined with silicone oil, ELR allows an aqueous solution to form spherical droplets with a 180-degree contact angle^18^. This ELR surface can be further modified by using a patterned stamp and oxygen plasma, removing parts of the ELR treatment and leaving a hydrophilic surface in the patterned shape. The result is a hydrophilic pattern on a repellant (superhydrophobic) background known as double-ELR^14^ (fig. 2a). For the application of bacterial isolation and expansion, a grid of five by eleven 2 mm spots is used to create an array of 2 µL droplets. This array allows for the combination of bacterial isolation and expansion by partitioning a concentrated bacterial sample into a large number of droplets, enabling an isolated expansion. Viable bacteria grown within the under-oil droplet array can be used for downstream molecular analysis methods such as microbe identification and antimicrobial susceptibility testing (AST). Distributing the aqueous sample within the array can be done in one of two ways. The first is called ‘sweep patterning’ or ‘sweep pipetting’, where the sample can be dispensed with a large bore pipette tip and manually dragged across the double-ELR pattern, leaving behind an equal portion of that sample as it moves (fig. 2b). The other method is standard pipetting onto each array spot using a 3D-printed patterning guide placed under the device for visualization of the deposited pattern (fig. 2c).

**Fig. 1.**
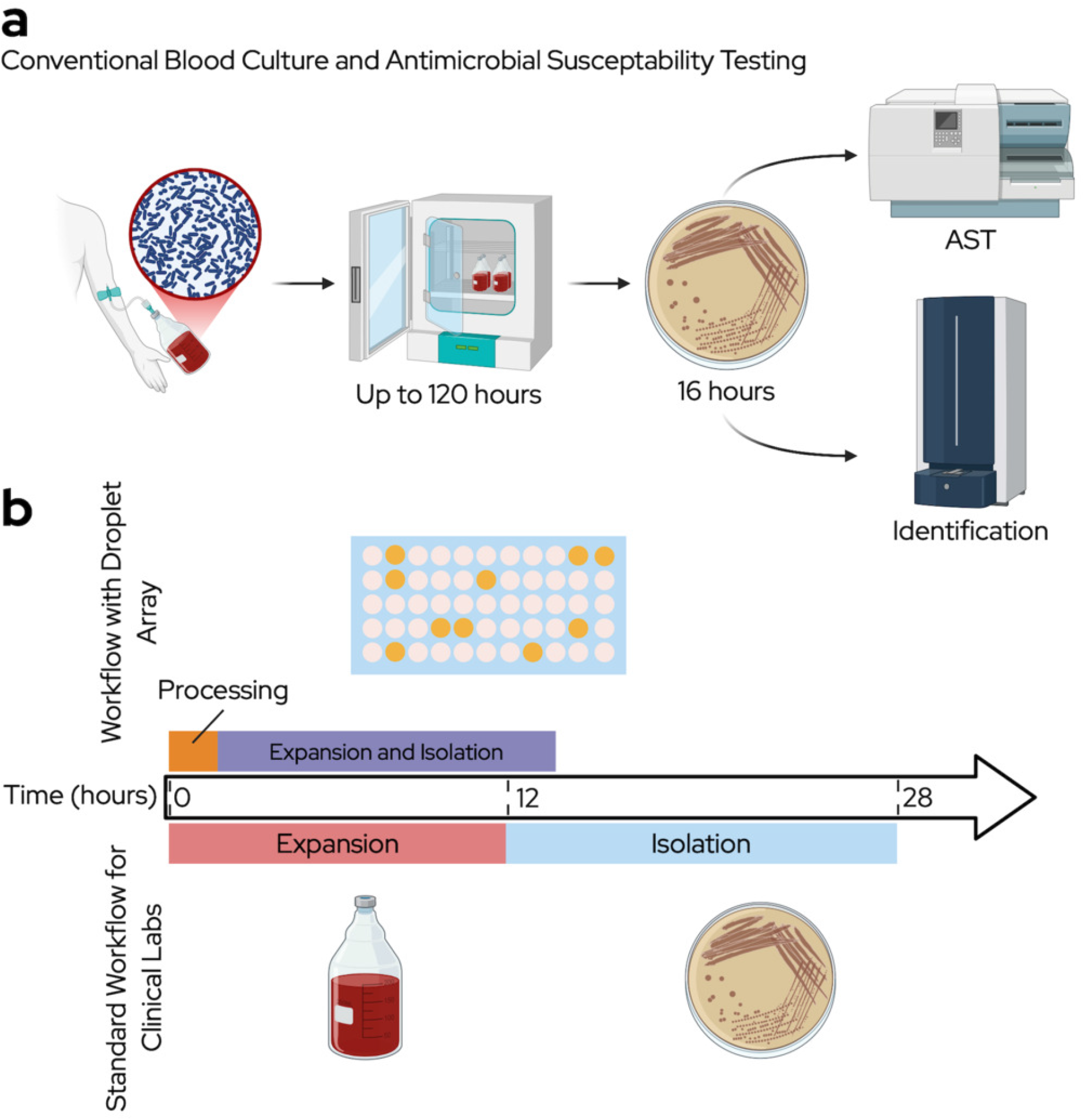
Overview of current laboratory workflow and proposed timeline. **a.** Schematic showing the current workflow utilized by clinical laboratories. **b**. Schematic depicting the timelines for both traditional workflow and the proposed droplet array workflow and the proposed under-oil droplet array workflow.

**Fig. 2.**
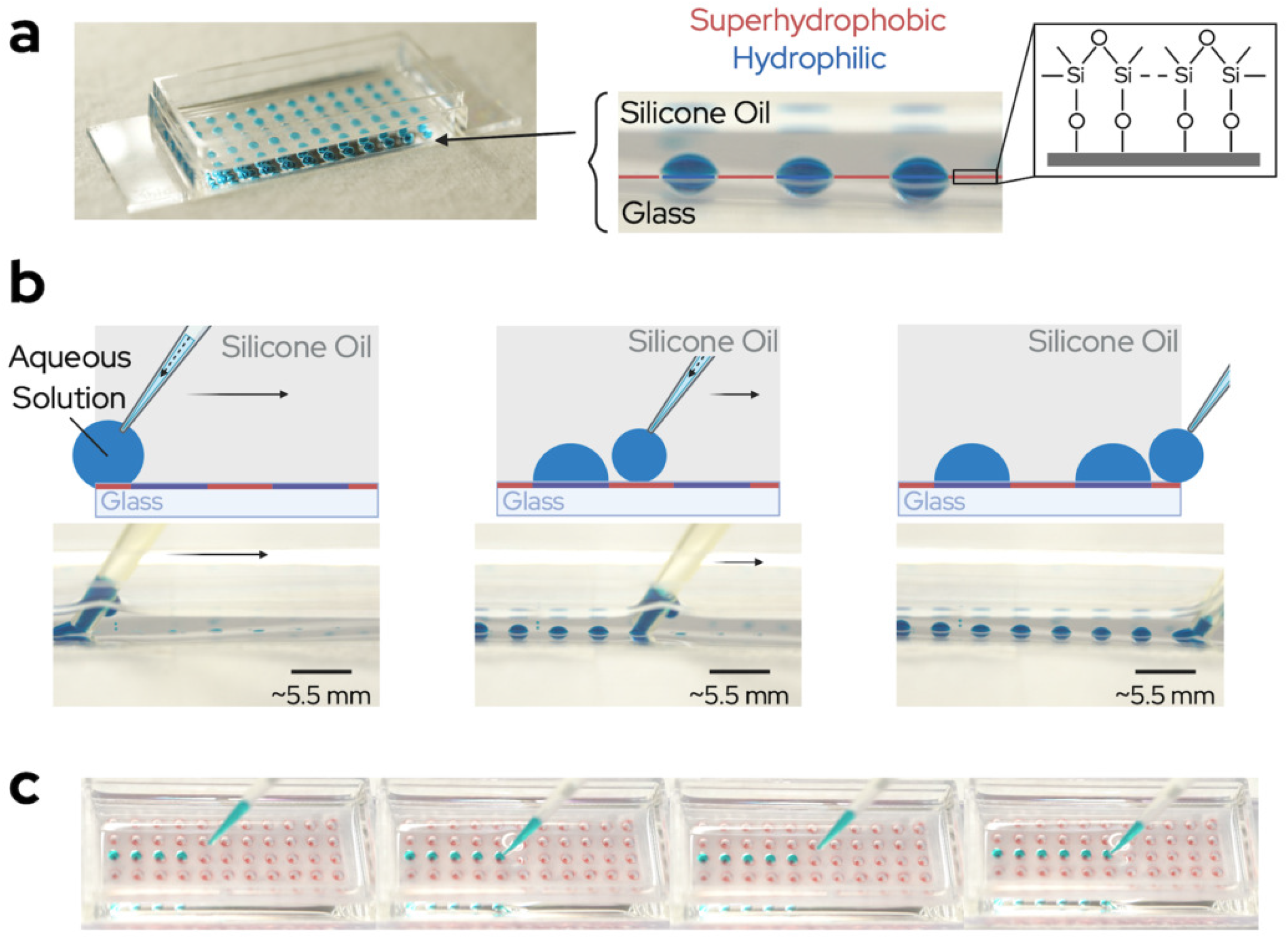
Features of double-ELR as an enabling technology. **a.** Visualization of ELR technology showing an under-oil droplet array consisting of a hydrophilic pattern on a superhydrophobic background with a silicon oil overlay. **b**. Diagram showing a unique method of pipetting enabled by ELR called sweep pipetting, where an aqueous sample can be deposited suing a continuous motion over the hydrophilic pattern. **c**. Images showing standard pipetting in the under-oil droplet array, made easier with an electronic repeater pipette and a guide that sits under the device, showing the location of the hydrophilic pattern.

### Blood processing for bacterial enrichment and whole blood clearance

To remove unwanted blood products and concentrate the bacterial sample into a workable volume, the blood sample must be processed. This workflow was developed specifically to use equipment and techniques utilized in most clinical laboratories. Blood processing consists of repeated centrifugation at 20,000 x g followed by targeted lysis using RBC lysis buffer or water (fig. 3a). Lysis works to neutralize whole blood products while leaving the bacteria unaffected, and centrifugation at high speed minimizes bacterial loss. The process leverages the biological differences between mammalian cells and bacteria to create a nonspecific bacterial isolation process that works for a wide range of bacteria, with this work specifically using seven different species, four gram negative and three gram positive. The ability to capture a wide range of bacteria is particularly valuable in a clinical setting where information about the potential pathogen may be limited. While sample processing resulted in a variable reduction in the number of viable bacteria, with a mean recovery of approximately 40%, all replicates yielded viable organisms, even at inoculum ≤5 CFU/mL (fig. 3b).

**Fig. 3.**
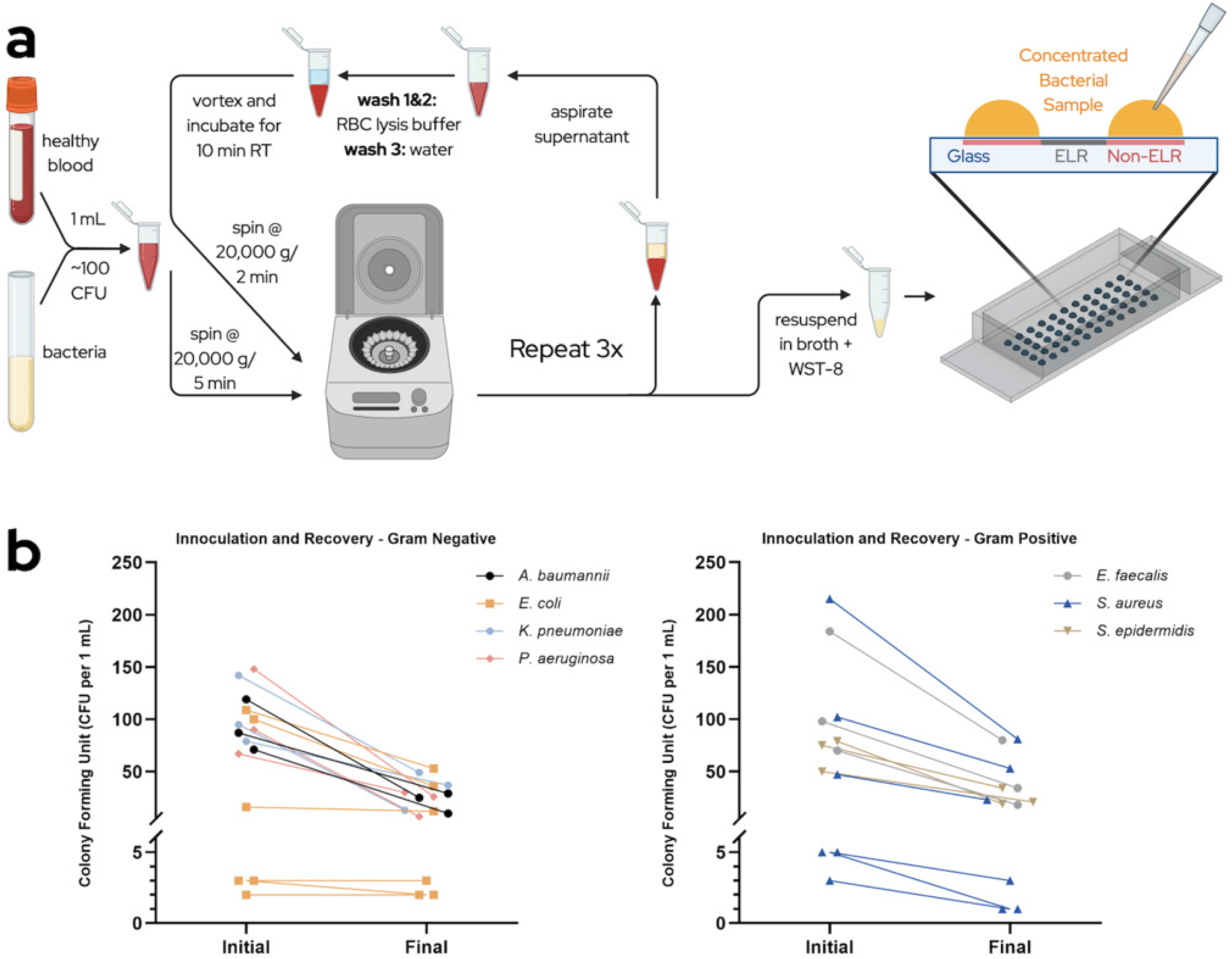
Blood processing and impacts on bacterial loads. **a.** Schematic of the steps used to process contrived blood samples and concentrate bacteria. **b**. Plots of initial bacterial concentrations compared to concentrations after blood processing. Initial bacteria bacterial loads were determined by counting the total colonies on an agar plate streaked in parallel with the same volume of bacteria spiked into blood samples. Final bacterial loads were determined by counting the total colonies on an agar plate streaked with a parallel sample after processing.

### Distribution of bacteria follows a Poisson distribution

Current clinical methods obtain single colonies of bacteria for analysis through agar subculture. The developed array also obtains single colonies of bacteria but leverages small volumes and Poisson distribution rather than solid growth medium. This means that at clinically relevant bacterial loads (∼100 CFU/mL), most droplets within the array will contain no bacterial growth, some droplets will contain one CFU, and even less contain two CFU.^16,17^ The probability that one CFU will be isolated decreases as the number of CFU per droplet increases. This can be modeled through the Poisson equation (equation 1).

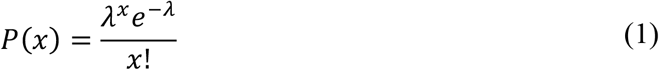

To model a standard Poisson distribution based on workflow specific data, the average rate (*λ*) for each replicate was determined (equation 2)

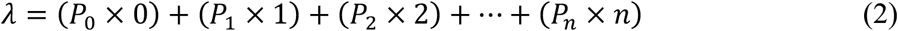

where *P*_*n*_ is the probability of *n* number of CFU in one droplet for that replicate. The average rate was then calculated for both gram-negative (*λ* = 0.288 ± 0.183) and gram-positive bacteria (*λ* = 0.320 ± 0.296). These rates were then used to calculate the standard Poisson distribution for gram-negative and gram-positive bacteria (fig. 4b). Of the droplets that contain any bacteria, the percentage of droplets containing a single CFU was calculated (equation 3)

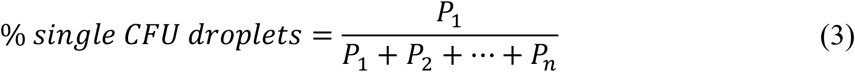

to be 86.2% for Gram-negative bacteria and 88.4% for Gram-positive.

**Fig. 4.**
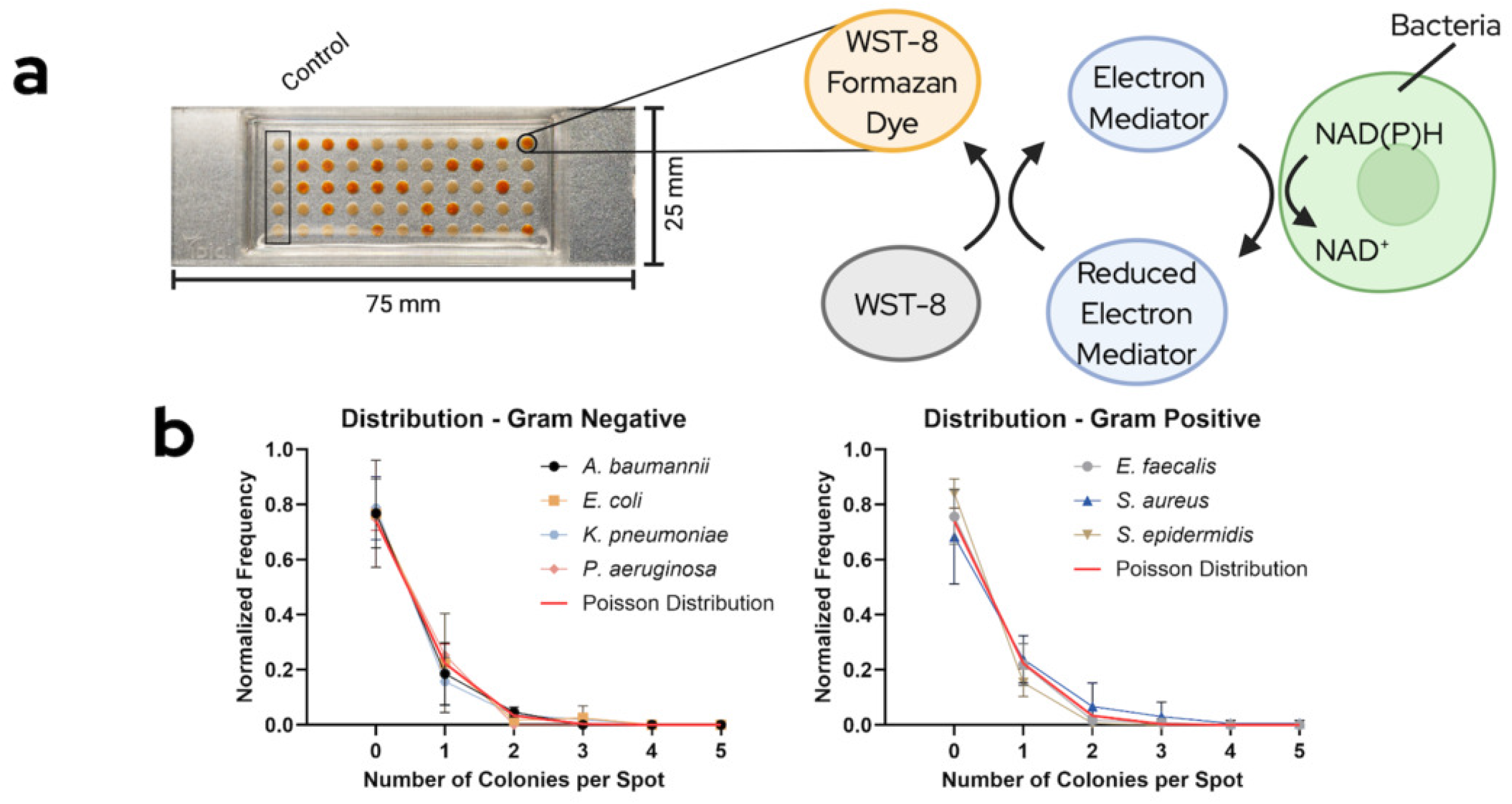
WST-8 dye and the distribution of bacteria in the Under-oil Droplet Array. **a.** Image of a device with S. aureus after 24 hours (contrived blood sample is concentrated in a volume of 100 µL, each droplet is 2µL for 50 droplets) and a schematic of how the WST-8 dye indicates bacterial growth. **b**. Distribution of colonies per droplet over different types of bacteria (n=3 per species).

### Colorimetric detection of bacterial growth and image thresholding

A colorimetric, visual indicator allows for visualization of bacterial growth in any specific droplet to be determined visually without the use of imaging systems or microscopy. WST-8 is a nearly colorless tetrazolium salt used to assess metabolic activity, converting NAD(P)H production at the bacterial cell membrane to water-soluble orange colored formazan dye (fig. 4a). WST-8 is commercially available and highly water soluble, forming a soluble formazan product rather than the insoluble precipitate caused by other formazan dyes.^19^ The amount of formazan produced is directly proportional to the number of viable bacteria and dependent on their metabolic activity. Blood processing steps were revised to use RBC lysis buffer after the WST-8 dye was selected to ensure a positive result (yellow/orange color) was more discernable (see “Material and Methods” section for additional details). Color thresholding analysis was performed via python script to objectively determine positive droplets and is a proof-of-concept for this devices compatibility with an automated detection system (fig. 5a,b). Color change for fast growing organisms such as *E. coli* and *K. pneumoniae* can be visually detected at around 10 hours, with *A. baumannii, S. aureus, P. aeruginosa*, and *S. epidermidis* being detectable closer to 12-14 hours (fig. 5c). While *E. faecalis* was successfully isolated and expanded in the droplets, it failed to react with the WST-8 when isolated from a contrived blood sample. A set value of yellow pixel fraction was used to determine a positive droplet (fig. 5d).

**Fig. 5.**
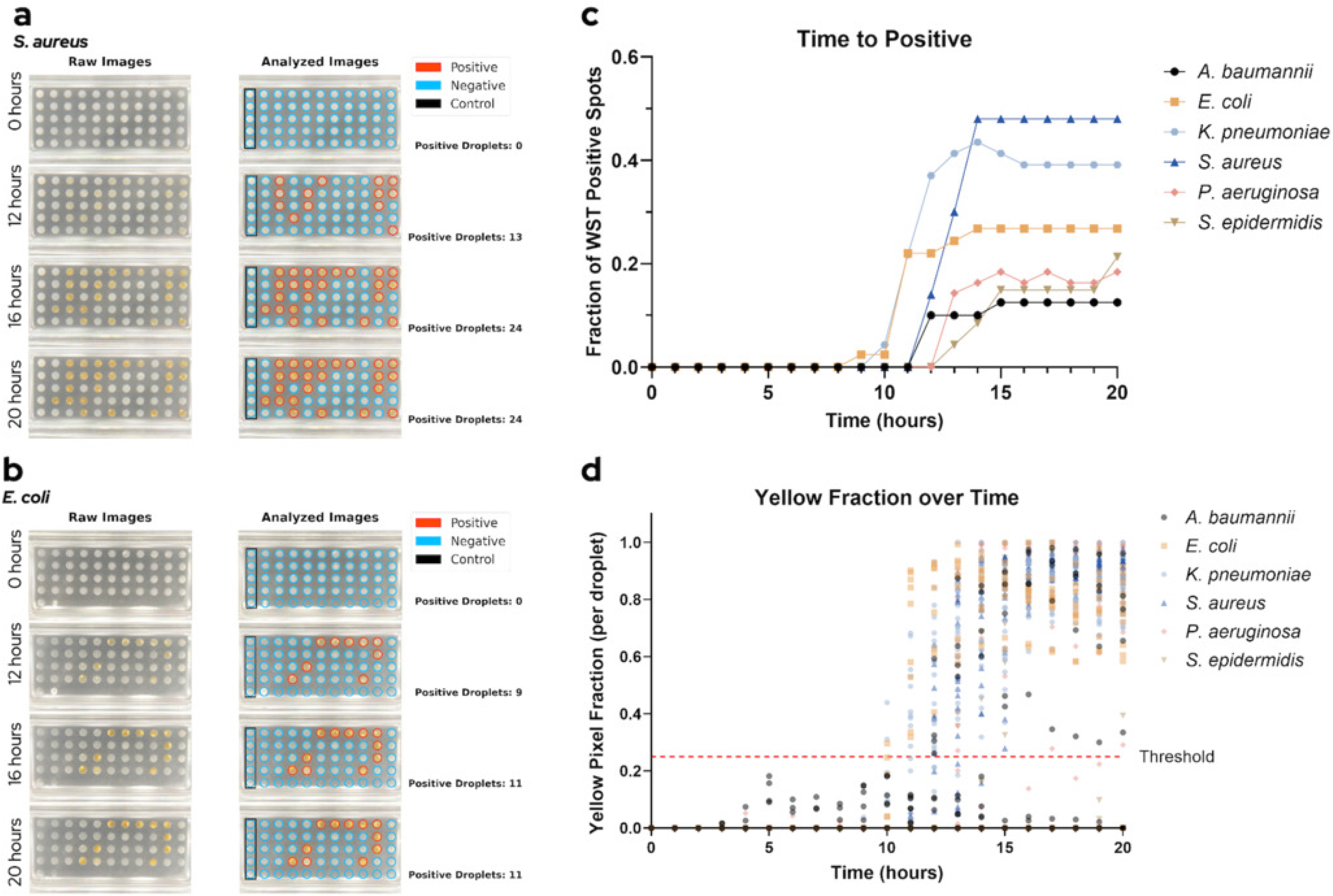
Time to detection of WST-8 dye and analysis. **a.** Methicillin-resistant S. aureus growth visible via WST-8 over time after isolation from whole blood, along with automated color detection with python script (see supplementary for python code). **b**. E. coli growth visible via WST-8 over time after isolation from whole blood, along with automated color detection with python script. **c**. Graph showing WST-8 positive droplets over time for all tested bacteria as detected by the python script. **d**. Graph of yellow pixel fraction of each droplet over time for all tested bacteria, showing the positive threshold at 0.25

Clinical samples can be more complex than a single strain of bacteria with a single form of resistance. Standard blood culture may lose some of this complexity in the incubation step through the process of competitive exclusion, where the fastest growing strain(s) dominates the culture. The described workflow utilizing blood processing and an under-oil droplet array allows for isolation and subsequent expansion of bacteria, increasing the likelihood of detecting multiple strains (polymicrobial infection) or heteroresistance within a single sample. To demonstrate the potential for the described workflow to enable detection of polymicrobial infection and heteroresistance, we inoculated blood samples with two species of labeled bacteria at relevant concentrations. Polymicrobial infection was simulated using equal parts *E. coli* and *S. aureus*, while heteroresistance was simulated using a 9:1 ratio of methicillin-susceptible *S. aureus* (MSSA) to methicillin-resistant *S. aureus* (MRSA) (fig. 6a,b). For polymicrobial simulation, growth occurred in individual wells for each species, with select wells displaying both species. Similarly,the small subpopulation of heteroresistance (MRSA) was isolated in select wells from the larger dominant population of MSSA. This partitioning has the added benefit of also preventing single or contaminating strains from dominating a culture.

**Fig. 6.**
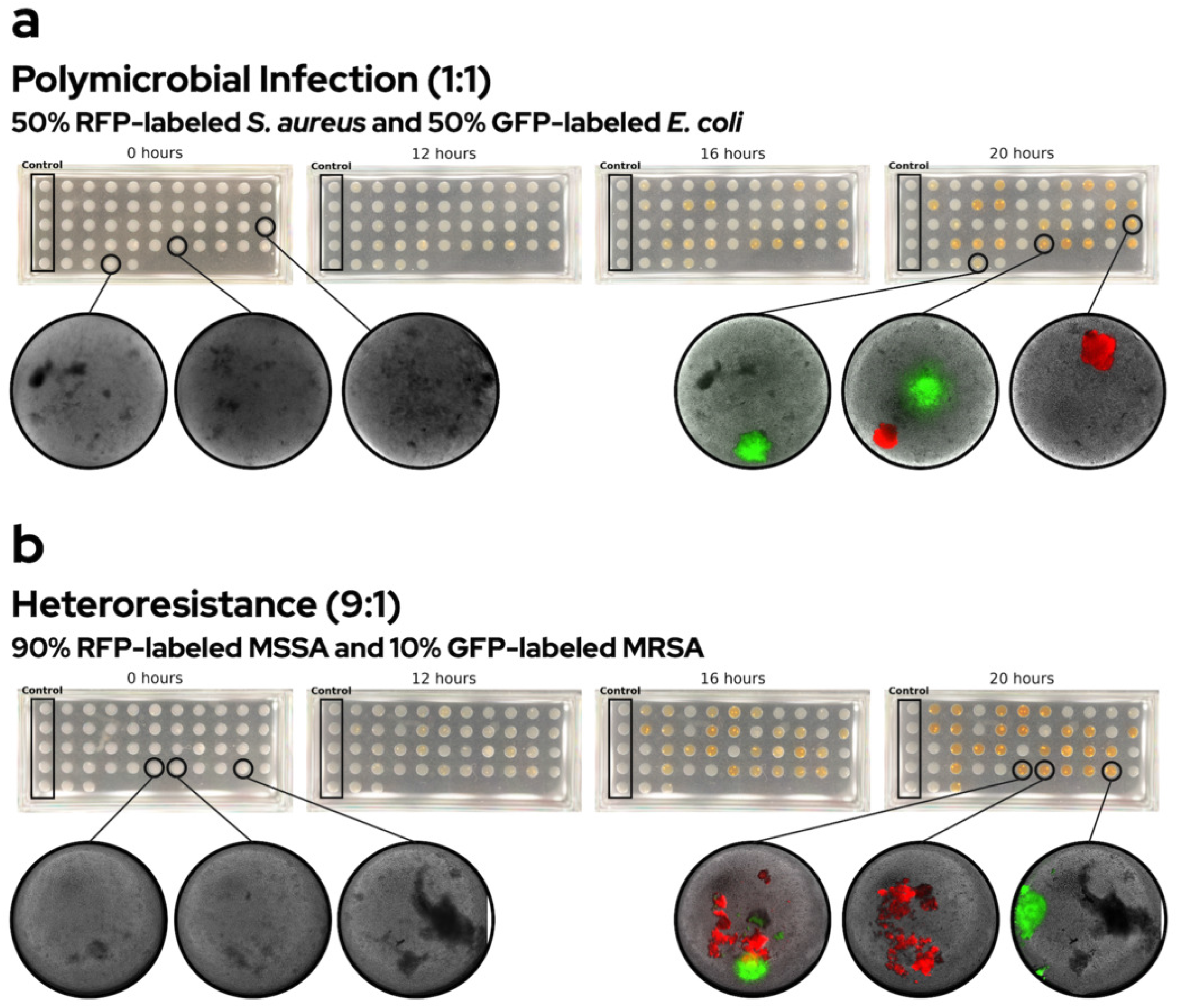
Utility in cases of polymicrobial infection and heteroresistance. **a.** Schematic showing how the device effectively isolates different bacteria in a sample simulating polymicrobial sepsis (equal inoculation of RFP-labeled S. aureus and GFP-labeled E. coli. **b**. Schematic showing how the device effectively isolates a resistant subpopulation of bacteria in a sample simulating heteroresistance (9:1 inoculation of RFP-MSSA and GFP-MRSA, respectively)

## Discussion

This study presents a novel methodological alternative to traditional blood culture, leveraging common laboratory equipment and a droplet array to isolate and expand bacteria in a single step. Here, we show that this device can reliably isolate 1 CFU per droplet over seven common sepsis-causing species (4 gram negative, 3 gram positive).

Combining blood culture and subculture into a single step provides unique advantages over traditional and existing blood culture methods. Primarily, a reduction in time by eliminating the need for agar subculture accelerates the overall process of resulting pathogen identification and AST, allowing clinicians to adjust or begin targeted antibiotics sooner. Similarly, the isolation of bacteria before expansion reduces any endogenous antimicrobial components that exist in whole blood and attempts to preserve all viable bacteria separately. In cases such as polymicrobial infection, heteroresistance, or potential contamination, this prevents one strain or species from outcompeting another during expansion, providing a more complete view of the pathogen(s) infecting the patient.

Traditional blood culture requires 8-10 mL per culture bottle for an adult patient, with 1-5 mL per culture bottle scaled by weight/age for neonates and pediatric patients.^20^ This study uses 1 mL of blood volume per contrived sample. Reducing blood draw volume would ease the burden on patients with lower total blood volume, such as children and neonates. Both blood processing and the droplet array could accommodate larger volumes up to ∼4.5 mL with minimal changes. However, this low-volume process cannot be directly compared to traditional blood culture at similar volumes as the whole blood components are lysed before bacterial expansion. This means that bacteria in the sample are no longer inhibited by leukocytes and other whole blood components that conventional blood culture can only partially mitigate through resin or charcoal within the growth supplement. Because of this, the sensitivity of the new workflow depends on the number of organisms sampled and on the portion that survive the isolation process to form detectable colonies, which may partially offset the reduced capture probability of a smaller draw volume with standard procedures.

The described workflow uses standard techniques and equipment found in most clinical laboratories. This new approach allows for easier implementation into most clinical settings and presents the possibility of automation and standardization using commercially available liquid handling systems and automated colorimetric detection for larger hospital labs. Alternatively, the described workflow could be utilized by laboratories that lack automated blood culture infrastructure and must send their blood cultures to a more complex facility, acting as a standalone replacement or an additional method of isolation and expansion. The use of WST-8 eliminates the need for an advanced microscope and specialized training to detect bacterial growth, allowing for growth to be easily visible. The retail cost per slide is comparable to the retail cost of a blood culture bottle ($10-15) and necessary reagents (cell culture-grade water, RBC lysis buffer, broth, WST-8) are inexpensive.^21^ If commercially available, the overall costs of the described device compared to that of traditional blood culture would likely be reduced per patient, as there is no investment needed for specialized equipment. An optimized version of this workflow could help supplement care where diagnostic infrastructure is weakest, such as low-income regions, remote facilities, or areas disrupted by natural disasters or outbreaks.

While this work shows the unique utility and potential of this system, it also has important limitations. While every bacteria type tested grew in the under-oil droplet array, *E. faecalis* did not react with WST-8 when isolated from whole blood despite growth visible through microscopy after incubation. Further development and optimization of this process could provide support for a wider range of bacterial and yeast organisms through specialized media and metabolic indicators, similar to that seen in blood culture bottles. The future use of direct patient samples in this workflow will help validate compared to the contrived samples shown above and the current clinical methodology. Finally, while using color imaging and determining yellow pixel fraction is an effective method to showcase potential automation and present data, variations in both lighting and dye coloration can affect how a droplet is classified. Human judgement is accurate and cost effective until a more robust system can be developed.

In summary, the described workflow improves on current blood culture methods by providing viable, isolated bacteria faster than traditional blood culture and limiting expansion culture, while preventing competitive exclusion and enabling the isolation of heteroresistance and polymicrobial infection. The isolated bacteria can then be utilized for further identification and analysis, providing medical teams with additional information to make treatment decisions, improving patient outcomes, and reducing AMR.

## Acknowledgments

We would like to thank Dr. Caroline Grunewald for her work creating the RFP-labeled MSSA used in this work. This study was funded by the University of Wisconsin Carbone Cancer Center Support Grant and the National Institutes of Health (NIH R01AI154940, NIH R01AI184593).

## Code Availability

Code is available on github via https://github.com/ammolenda/DropletArrayCulture-Analysis.git

## Author Contributions

A.M., M.L., N.H., designed and performed experiments. B.H. assisted with understanding the clinical process and testing the protocol. S.M. assisted in microbial culture. W.E.R. and D.J.B. oversaw the study. A.M. and M.L. wrote the manuscript with input from all authors.

## Conflicts of Interest

David J. Beebe holds equity in Bellbrook Labs LLC, Salus Discovery LLC, Lynx Biosciences Inc., Stacks to the Future LLC, Flambeau Diagnostics LLC, Navitro Biosciences LLC, Eolas Diagnostics, Inc., and Onexio Biosystems LLC. All other authors declare no competing interests.

